# Genome-wide association study and regional heritability mapping of protein efficiency and performance traits in Swiss Large White pigs

**DOI:** 10.1101/2023.11.28.568963

**Authors:** Esther Oluwada Ewaoluwagbemiga, Audald Lloret-Villas, Adéla Nosková, Hubert Pausch, Claudia Kasper

## Abstract

**Background:** The improvement of protein efficiency (PE) is a key factor for a sustainable pig production as nitrogen excretion contributes substantially to environmental pollution. Protein efficiency has been shown to be clearly heritable and genetically correlated with some performance traits, such as feed conversion ratio (FCR) and average daily feed intake (ADFI). This study aimed to identify genomic regions associated with these traits through genome-wide association studies (GWAS) and regional heritability mapping (RHM) using whole-genome sequence variants from low-pass sequencing of more than 1,000 Swiss Large White pigs.

**Results:** The genomic-based heritability estimates using ∼15 million variants were moderate to high, ranging from 0.33 to 0.47. Using GWAS, no significant variants were found at the genome-wide thresholds for PE and FCR, but 45 variants were identified for ADFI on chromosome 1and one for ADG on chromosome 14. No region was found to be significantly associated with PE and FCR using RHM. We identified five suggestive regions on chromosome 1 for ADFI and one on chromosome 14 for ADG. Combining both analyses, we were able to highlight putative candidate genes for PE, which included *PHYKPL, COL23A1, PPFIBP2, GVIN1, SYT9, RBMXL2, ZNF215* and olfactory receptor genes.

**Conclusions:** Combining GWAS and RHM allowed us to suggest genomic regions that potentially influence PE and production traits. The apparent difficulty in detecting significant regions for these traits probably reflects the relatively small sample size, differences in genetic architecture across experimental conditions, and the possibility that polymorphisms explaining large parts of the trait variation may not segregate in the population. Nevertheless, we identified plausible functional candidate genes in the highlighted regions, including those involved in nutrient sensing, the urea cycle, and metabolic pathways, in particular IGF1-insulin), and have previously been reported in association with nitrogen metabolism in cattle, muscle and adipose tissue metabolism and feed intake in pigs. We also highlighted a range of noncoding RNAs, and their targets and gene regulation in general in this context should be investigated in the future.

## Background

Efficient livestock production is gaining importance due to the increasing global demand for meat that has led to an increased environmental pollution. A key pollutant in livestock production is nitrogen, which forms harmful compounds such as nitrate, ammonia and nitrous oxide [1, 2, 3]. Although pigs are more protein efficient than cattle [4], with nitrogen excretion of about 70-80% in beef and dairy production [5, 6, 7], approximately 50% of the dietary protein consumed by pigs is excreted as waste [8, 9]. Undigested proteins are excreted in faeces, and absorbed amino acids not used for growth or maintenance are eliminated in urine [10]. This is an environmental concern because the nitrogen in urine is in a volatile form [11, 12]. The percentage of nitrogen in urine ranges from 48.6 % on a pectin-enriched diet [12] to 79.3 % on a grain-based diet [11] and is influenced by the fibre content of the diet, its fermentability, nitrogen digestibility and the pig’s growth and maintenance requirements. Methods such as reducing dietary nitrogen [13, 14] and selection to increase protein efficiency (PE; the proportion of total dietary protein intake retained in the carcass) in pigs [10] have been proposed to reduce the contribution of animal-based food production to environmental pollution. Varying heritability estimates between 0.14 and 0.59 have been recently reported for PE and related traits (e.g., nitrogen digestible coefficient, protein deposition), depending on the breed, fattening phase and diet type [15]. The significant genetic variance and moderate to high heritability (h^2^) estimates for PE and nitrogen use efficiency from these studies (ranging from 0.16 to 0.54) [15] indicate that this trait can be genetically improved and thus presents a promising target towards a more sustainable pig production through reduced nitrogen excretion. Performance traits such as feed conversion ratio (FCR), average daily feed intake (ADFI), and average daily gain (ADG) are also important, considering their economic and environmental impacts. Several studies have reported moderate heritabilities for these traits [10, 16] and genetic correlations among them in pigs [15]. Genetic correlations (± SE) in Large White pigs of -0.55 ± 0.14 between PE and FCR [10], digestibility coefficient of nitrogen (DCN) and FCR of -0.24 ± 0.10 [17], -0.53 ± 0.14 between PE and ADFI [10], DCN and ADFI of -0.53 ± 0.13 [17], -0.19 ± 0.19 between PE and ADG [10] and -0.15 ± 0.17 between DCN and ADG [17] have been previously reported. There was also a strong negative genetic correlation between the digestibility of protein (DP) and the FCR in broiler chicken [18].

Genome-wide association studies (GWASs) in pigs have reported loci associated with several important traits, such as meat quality [19], performance [20, 21], carcass [22], body composition [23], and efficiency-related traits [24]. However, despite the environmental importance of nutrient efficiency traits, such as PE and nitrogen excretion, to date, only the studies of Shirali et al. [25] and Schmid et al. [26] have identified genomic regions associated with nitrogen-excretion traits and nitrogen use efficiency, respectively, in pigs. The study of Shirali et al. [25] used 315 pigs from Piétrain grand-sires and grand-dams from a three-way cross, with pigs genotyped for 88 microsatellite markers on 10 (of 18) chromosomes [25]. Their study identified three quantitative trait loci (QTL) associated with total nitrogen excretion throughout the 60 – 140 kg live body weight (BW) growth period on chromosomes (SSC) 2, 4, and 7 [25]. Three additional QTL were found for another excretion trait – average daily nitrogen excretion – on SSC 6, 9, and 14 for the same growth period [25]. However, the study by Shirali et al. [25] is limited by its very small sample size and the use of a small number of markers that are not evenly distributed across the genome. Schmid et al. [26] reported significant SNPs for nitrogen use efficiency in German Landrace×Piétrain crosses on SSC 5 and 13 during fattening phase 1 (40kg BW) and on SSC 6 only during fattening phase 2 (60kg BW) at a significance level of 5 × 10^−5^. Although the study included a considerably higher number of markers (over 48,000 SNPs), the sample size was relatively limited (N=502). Furthermore, the nitrogen use efficiency phenotype was predicted from N-balance data and blood metabolites measured in only 10% of the animals [26]. It is also noteworthy that both studies involved Piétrain crossbreeds, which are likely to differ significantly in genetic background from the animals in this study (see [15] for discussion).

As FCR, ADG, and ADFI have either direct or indirect impacts on efficiency and production costs, a number of studies have identified QTL for FCR, ADG and ADFI [26, 27, 28, 29, 30]. The majority of these QTL were found in Duroc and Landrace pigs, with only a few QTL identified in Large White pigs [31]. However, although FCR and RFI may be correlated with PE as reported in the studies by Ewaoluwagbemiga et al. [10] and Saintilan et al. [16], it has been suggested that selection for improved FCR and RFI with the aim of reducing nutrient excretion is clearly less efficient than direct selection for the nutrient efficiency trait itself (e.g., PE) in poultry [32].

Complex traits are typically influenced by many genes (i.e., are polygenic), with many genetic variants having too small effect sizes to be detected at the Bonferroni-corrected or false-discovery rate (FDR) threshold of GWAS [33, 34]. This makes it difficult to identify associated variants, particularly for traits for which limited sample sizes are available. Besides GWAS, regional heritability mapping (RHM) offers an alternative approach to identify genotype–phenotype associations [35, 36]. RHM divides the genome into windows of a certain number of variants. Subsequently, a genomic relationship matrix (GRM) is computed using all the variants in each region, and the variance of the trait explained by each region is estimated [37]. RHM is used to identify regions that contribute to the genetic variance of a trait by aggregating the effects of multiple variants within a defined region, rather than analyzing them individually [37]. This approach can potentially capture signals from variants with effects too small to be detected by single-SNP GWAS, albeit at the cost of lower resolution because it estimates the heritability of broader genomic regions rather than pinpointing specific variants. By grouping multiple variants, RHM reduces the number of statistical tests required, which may allow for less stringent correction for multiple testing. However, RHM’s power to detect associations depends on the extent to which genetic variation within the region is collectively influential, and its performance may vary with factors such as regional linkage disequilibrium structure and trait architecture.

A limitation of RHM is that it estimates the contribution of a region to trait variance by modeling it as a random effect, which generally requires greater statistical power (i.e., larger sample sizes) to obtain reliable estimates. This is in contrast to GWAS, which typically estimates fixed effects for individual variants and thus involves simpler parameter estimation. As a result, RHM may be less efficient in smaller datasets or when regional effects are modest.The RHM approach has been shown to be effective in other studies; for example, Resende et al. [38] detected 26 QTL associated with 7 traits in eucalyptus using RHM, compared to only 13 QTL identified by GWAS. Similarly, Sutera et al [39] found 5 QTL associated with fat percentage in sheep using RHM. In this study, we used RHM alongside GWAS to strengthen the evidence for suggestive variants with the aim of providing a complementary analysis that can validate and clarify results that may otherwise remain inconclusive.

The aim of this study was therefore to investigate the genetic basis of PE and performance traits in Swiss Large White pigs. This was achieved by estimating the genomic heritability and performing GWAS and RHM using low-pass sequence data.

## Methods

### Animals and phenotypes

We analysed a total of 1,036 Swiss Large White pigs, which were previously included in several nutrition experiments and one genetic study. The Swiss Large White dam line used in our study is selected for high fertility with good piglet rearing ability, but also, albeit to a lesser extent than the sire line, for production. The relative weighting of fattening performance, feed conversion and lean meat content together account for around 30 % of the selection index and meat quality accounts for around 14 % of the selection index). To produce crossbred fattening pigs in practice, a crossbred sow (Large White dam line × Swiss Landrace) and a boar from the Large White sire line are used, the latter being primarily selected for meat quality, fattening performance, FCR and lean meat content (37%, 26 %, 19 % and 8 % relative weighting in the selectin index, respectively). The data set used in this study is described in detail in Ewaoluwagbemiga et al. [10]. Briefly, data from 7 experiments, which were carried out at Agroscope Posieux in Switzerland, were combined. Pigs had *ad libitum* access to isocaloric diets that differed in crude protein or fibre content. The majority of pigs (853) received a diet that contained 80% of the crude protein and digestible essential amino acids content of the control diets in both grower and finisher phase, and 16 pigs received a standard (control) diet in the grower and the protein-reduced diet only in the finisher phase. The three control groups had no reduction in dietary crude protein, and diets were mainly formulated according to the Swiss feeding recommendations for pigs^1^. One control diet (154 pigs) conformed to the recommendations in all nutrients. Two control diets were used in an organic pig production trial and contained 95 % to 100 % organic ingredients (32 pigs). A third 100% organic control diet was formulated to contain 6 % more crude fibres than the other controls (16 pigs). In all experiments, pigs were fed a grower diet from approximately 20 to 60 kg live BW and a finisher diet from 60 kg to slaughter at 100 kg. Most pigs (840) were slaughtered at about 100 kg BW, 32 pigs at 40 kg, 57 at 60 kg, 50 at 80 kg, 53 at 120 kg and 40 at 140 kg live BW. Pigs that were slaughtered at 120 and 140 kg were fed another, specially formulated, finisher diet from 100 to 140 kg [40]. Most of the animals were females (492) or castrated males (469), only 110 were males. To account for a part of the heterogeneity in the data, we included the diet type (i.e. treatment), slaughter weight and sex as fixed effects in the model.

Every week, pigs were weighed individually, and, once a pig reached a live BW of approximately 20 kg, it was allocated to grower-finisher pens and the experimental treatments were started. This was done until a maximum number of 12 (or 24 or 48) pigs per pen (depending on the pen layout; minimum 1m^2^ per pig and maximum 12 pigs/feeder) was reached. Pigs remained in their pen until slaughter. Piglets were weaned at an average age of 27 ± 2 days after birth by removing the sow and were fed a standard starter diet with crude protein levels following the recommendation. At 22.3 (± 1.6) kg, pigs were placed in pens equipped with automatic feeders (single-spaced automatic feeder stations with individual pig recognition system by Schauer Maschinenfabrik GmbH & Co. KG, Prambachkirchen, Austria) and stayed in this pen until slaughter. The automatic feeder recorded all visits and feed consumption per visit, from which the total feed intake of each pig was calculated. The protein content of feed was monitored during production by near-infrared spectroscopy for each 500 kg batch. To obtain more accurate data on feed composition at the time of consumption, a sample was taken from each automatic feeder station each week, and the crude protein content was determined by wet-chemistry methods.

### Phenotype data

The phenotypes were derived as reported in Ewaoluwagbemiga et al. [10]. Total and average daily feed intake (ADFI) were recorded, and average daily gain (ADG) and the feed conversion ratio (FCR) were calculated as follows:

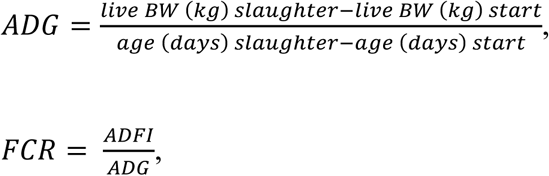

where *live BW* (*kg*) *slaughter* and *age* (*days*) *slaughter* are the live pre-slaughter body weight in kg and the age in days at slaughter, respectively, and *live BW* (*kg*) *start* and *age* (*days*) *start* are the exact body weight in kg and the age in days at the start of the grower phase, respectively. To measure PE, the left carcass half of each pig, including the whole head and tail, was scanned with a dual-energy X-ray absorptiometry (DXA; GE Lunar i-DXA, GE Medical Systems, Glattbrugg, Switzerland) to determine the lean tissue content, which was used in the equation of Kasper et al. [41] to estimate the protein content retained in the carcass;

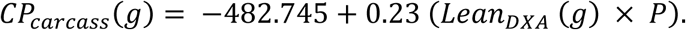

Where *CP*_*carcass*_(*g*) is the crude protein content of the carcass in g, *Lean*_*DXA*_ (*g*) is the lean meat content obtained with DXA in g, and *P* is the proportion of the left cold carcass (also includes the whole head and tail) to the total cold carcass weight. This method of estimating carcass protein content using DXA yields a highly precise and accurate phenotype with an R^2^ between 0.983 – 0.998 [41, 42]. Unlike other techniques, such as estimating carcass protein using semi-mechanistic models [43] or determining protein digestibility from faeces with near-infrared spectroscopy [17], This approach allows calculating PE as protein retained over protein intake that does not rely on assumptions and is therefore likely to preserve individual differences in the trait. PE was calculated as;

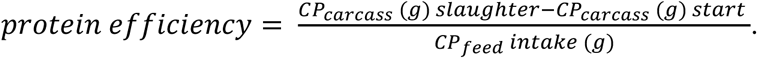

Where *CP*_*carcass*_ (*g*) *slaughter* is the crude protein content of carcass at slaughter, *CP*_*carcass*_ (*g*) *start* is the crude protein content of carcass at the start of the experiment, and *CP*_*feed*_ *intake* (*g*) is the crude protein feed intake. The crude protein content of pigs at the start of this experiment (*CP*_*carcass*_ (*g*)*start*) was estimated from a sample of 38 piglets (12 females, 12 castrated males and 14 entire males). These piglets were slaughtered at an average of 20.98 ± 1.85 kg BW in a previous experiment, and their carcass protein content was chemically determined [40]. The average protein content per kg carcass for each sex (female, entire male, castrated male) was used to estimate *CP*_*carcass*_ (*g*) *start* for the pigs by multiplying the actual live BW of pigs when they entered the experiment (i.e., at approximately 20 kg body weight) with the protein content per kg carcass of piglet, as previously determined from the 38 piglets [40].

### Genotype data and imputation

DNA was extracted from blood, and the sampled pigs were genotyped on three different platforms, namely the Affymetrix 600K axiom porcine genotyping array, whole-genome sequence data at an intended 4-fold coverage, and low-pass sequence data at an intended 1-fold coverage with Gencove. Thus, the three genotyping/sequencing platforms comprised of (i) 258 pigs genotyped at 600K obtained with the Axiom Porcine Genotyping array; (ii) 297 pigs sequenced at an intended read depth of 4×; and (iii) 492 pigs sequenced at an intended read depth of 1×.

#### Imputation of array data

The array genotyping data was imputed to whole genome sequence level with a reference panel consisting of 421 pigs (Landrace and Large White, including the 297 pigs in (ii)) that were sequenced at a coverage ranging between 4× and 37.5× using Beagle (v 4.1) [44], and had imputation accuracy of 0.92 (R^2^) [31, 44]. The imputed array data finally contained 29,469,425 autosomal variants (SNPs and indels).

#### Filtering and imputation of 4× sequencing data

We estimated the average realized coverage as 4.5× ± 0.9× (calculated as the amount of raw data obtained divided by the approximate pig genome size of 2.8 GB). Filtering and alignment to the reference genome were carried out as described in [45]. In brief, raw reads with a phred quality score below 15 for more than 15% of the bases were removed using the the fastp software (version 0.19.4) with default parameter settings [46]. The filtered reads were aligned to the Sus Scrofa 11.1 assembly [47] using the MEM-algorithm of the Burrows-Wheeler Alignment (BWA) software (version 0.7.17) [48, 49]. Subsequently, the alignment quality and coverage depth were assessed. Duplicate reads and reads with a mapping quality ≤ 10 were removed. The imputation of sporadically missing genotypes in the variant files was done using Beagle (v 4.1) [44] with an imputation accuracy of 0.98. The 4× sequenced data contained 30,179,303 variants.

#### Filtering and imputation of 1× sequencing data

The 1× data was imputed by Gencove’s loimpute pipeline v0.1.5 [50] using 414 publicly available pig sequence data with an imputation accuracy of 0.97. The data contained 45,100,556 autosomal variants (including 13,361,070 non-variant sites) and the realized coverage after mapping and variant calling was 0.61×.

#### Merged variant call format file

Eight pigs without phenotypes and three pigs with a mis-match between pedigree and genomic-based relationship matrix were excluded from the further analyses. PLINK (v1.9) [51, 52] was used to merge the three different SNP panels based on their physical positions according to the reference genome (*Sus Scrofa 11.1*). After merging, there were 23,171,650 intersecting biallelic variants (including indels) and 1,036 individuals.

### Genome-wide association study

Prior to GWAS, we tested for outliers in the phenotypes, and removed individuals with phenotypes not in the range of µ ± 3σ. This resulted in 1025, 1033, 1034, and 1024 individuals remaining for PE, ADG, ADFI, and FCR, respectively. For each trait, we removed variants with minor allele frequency (MAF) < 5% and variants that deviated from Hardy-Weinberg equilibrium (P < 0.0001). After quality control, 15,269,953, 15,192,400, 15,200,584, and 15,220,328 variants were included for PE, ADFI, ADG, and FCR, respectively.

The residuals of linear mixed-effects models in R software v 4.2.1 [53, 54] adjusting for environmental effects were used as phenotypes in GWAS for each trait. The environmental effects included the fixed effects from a model selection step prior to estimating genetic parameters as described in Ewaoluwagbemiga et al. [10]. In brief, the fixed effects included year (factor variable), dietary treatment (factor variable), sex (factor variable; castrates, females and males), slaughter weight, ambient temperature in the barn at the start of the experiment, slaughter age, interaction of slaughter weight and sex, interaction of treatment and sex, interaction of treatment and slaughter age, and interaction of year and slaughter age. However, it should be noted that while the model accounts for these fixed effects, the genetic architecture of the traits may vary between different conditions, potentially reducing the statistical power to detect associated variants and leading to biased or inaccurate estimates of variant effects.

The GWAS was performed with GCTA using the fastGWA method [55], where SNP effects were tested using a linear mixed effects model approach, incorporating the genomic relationship matrix (GRM) to account for relatedness in the sampled population. The linear mixed effects model fitted to the data was

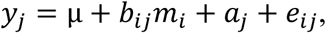

where *y*_*j*_ is a vector of residuals of phenotypes corrected for environmental effects; µ is the overall mean; *b*_*ij*_ are the marker genotypes, coded as 0, 1, and 2, of the *i^th^* variant for the *j^th^* individual; *m*_*i*_ is the allele substitution effect of the *i^th^* variant; *a*_*j*_ is the random polygenic effect of the *j^th^* individual following the distribution 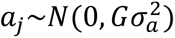, *G* is the GRM and 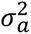 is the additive genetic variance; *e_ij_* is the random residual effect with 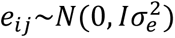, *I* is an identity matrix and 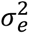 is the residual variance.

We applied a twofold approach for obtaining meaningful significance thresholds for GWAS that balances the need to correct for multiple testing and the fact that several variants tested are in strong LD and thus not independent. First, we used an empirical procedure to obtain a distribution of the test statistic (in this case the p-value) under the null hypothesis. To do so, we followed the procedure for permutation testing described in van den Berg et al. [56], which was based on Churchill and Doerge [57]. Specifically, we ran 1,000 GWASs for each of the traits using the same model and the same method as described above, but with the phenotype vector (i.e. the residuals) randomly shuffled in each run. For each iteration, we recorded the smallest p-value and we defined the experiment-wise critical value as the -log_10_ of the p-value at the 95th percentile [57]. Second, we applied a Bonferroni-corrected significance threshold at an alpha level of 0.05, based on the number of sufficiently independent loci derived from the typical LD-decay. In commercial pig lines, markers at distances of 100 to 150 kb have been shown to have an average r^2^ of about 0.3 [58]. Based on this, we chose a slighltly more conservative approach and pruned the data using Plink 1.9 (- - indep-pairwise 100 10 0.4) and used the number of remaining variants to calculate the threshold. We inspected the quantile-quantile (QQ) plots for the inflation of small p-values, which could indicate population structure. The genomic inflation factor was used to investigate a potential violation of the distribution assumptions of non-associated variants, which can arise due to population stratification [59]. Genomic inflation was calculated as the median of the observed chi-squared test statistics divided by the distribution of the expected chi-square test statistics.

### Regional heritability mapping

Regional heritability mapping was performed using GCTA software [55]. For this analysis, each chromosome (SSC1 to SSC18) was divided into several windows that contained 5,000 variants-each and that overlapped by 2,500 variants.Windows at the end of the chromosomes contained on average 3,837.5 variants, but never less than 2,533. Subsequently, the genomic variance was estimated for each of the around 5,230 regions of the whole genome (their exact number depended on the trait). The linear mixed effects model below was used to test the effect of all variants within each genomic region, which included the random regional genomic effect and the random genomic effect of the rest of the genome, including the specific region:

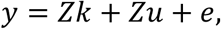

where *y* is a vector of the residuals of phenotypes corrected for environmental effects as indicated above for the GWAS; *Z* is the design matrix for the random effects; *k* is the random regional additive genomic effect following the distribution 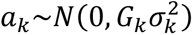 and *u* is the random polygenic effect of all variants following the distribution 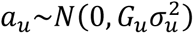. *G* is the regional GRM, 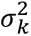 is the regional variance, *G*_*u*_ is the genome GRM and 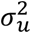 is the genome additive genetic variance. Regional and genome heritability were estimated as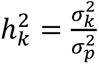 and 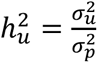, respectively, where 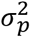 is the sum of the regional variance 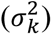, genome variance 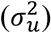, and residual variance 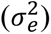. The statistical significance of the variance of a region was tested using the likelihood ratio test (LRT), which compares the log likelihood of the full model (including regional and genome variance) with the reduced model (including only genome variance). This was done by specifying the –reml-lrt 1 option in GCTA, which gives the LRT and p-value of the random regional additive genomic effect.

To identify significant and suggestive variants, two different thresholds were defined. It should be noted that the thresholds in addition to the Bonferroni correction differ between GWAS and RHM in our study. For the genome-wide significance threshold (Bonferroni correction for multiple testing) was applied at an alpha level of 0.05 for the around 5,230 regions, divided by 2 to account for the overlap. Instead of a LD-pruned threshold as used for GWAS, we set a suggestive threshold for RHM following the procedure described in [39]. Briefly, the suggestive threshold implies that, at every genome scan, one false positive is expected [39]. The thresholds applied in the current study, for RHM, were thus at p-values of 1.94×10^−5^ (-log_10_(p) = 4.71) and 3.89×10^−4^ (-log_10_(p) = 3.413) for the genome-wide 5% significance and the suggestive threshold, respectively.

### Identification of genes

After performing GWAS and RHM, we identified genes located within the significant and/or suggestive regions. These genes were identified using the pig reference genome (*Sus Scrofa 11.1*) on the genome data viewer^2^ [60], using the NCBI Sus scrofa Annotation Release 106 and Ensembl release 110 (Accession NC_010444.4). The biological functions of candidate genes were found with Gene Ontology database. The platforms AnimalQTLdb [61] pigGTEx^3^ [62] were used to retrieve additional information on previously identified QTL and molecular QTL, such as expression QTL (eQTL), QTL for long noncoding RNA expression (lncQTL), enhancer QTL (enQTL) and alternative splicing QTL (sQTL). We restrict reporting to tissues of interest for our traits, such as muscle, liver, tissues of the immune system and the gastrointestinal tract, and the brain.

## Results

### GWAS

The Bonferroni threshold based on LD pruned data for all traits, at the alpha level of 0.05, was 9.90 × 10^−8^, 9.93 × 10^−8^, 9.94 × 10^−8^, 9.91 × 10^−8^ for PE, ADG, ADFI and FCR, respectively. The threshold obtained with the permutation approach was 4.47 × 10^−8^, 3.56 × 10^−8^, 3.19 × 10^−8^, and 3.93 × 10^−8^, respectively, for PE, ADG, ADFI and FCR. No significant variants were found at both thresholds for PE (Figure 1, Table 1, Table S1). For ADG, 1 variant on SSC14 passed the LD-pruned threshold (Figure 2, Table 2, Table S2), for ADFI, 26 variants on SSC1 (25 SNPs and 1 indel) passed the permutation threshold and 19 further variants on SSC1 (15 SNPs and 4 indels) also passed the LD-pruned threshold (Figure 3, Table 3, Table S3). No variants passed either threshold for FCR (Figure 4, Table 4, Table S4). The heritability (±SE) of the suggestive variant for ADG was h^2^ = 0.02 ± 0.03, around 7% of the total genomic heritability, and the heritability of the 45 variants for ADFI together was h^2^ = 0.06 ± 0.04, around 14% of the total genomic heritability . The QQ plots of the GWAS analyses (Figure 5A-D) and the genomic inflation factor, which was approximately 1 for all the traits (PE = 1.08, ADG = 1.06, ADFI = 1.13, FCR = 1.05), suggested that population stratification has been sufficiently accounted for by the GRM. Although no or few suggestive associations were found for PE and FCR, the genomic heritability for all traits ranged from 0.33 to 0.47 using all available variants for each trait (Table 5), with ADG having a slightly higher genomic heritability (0.47) than pedigree-based heritability (0.45).

**Figure 1.**
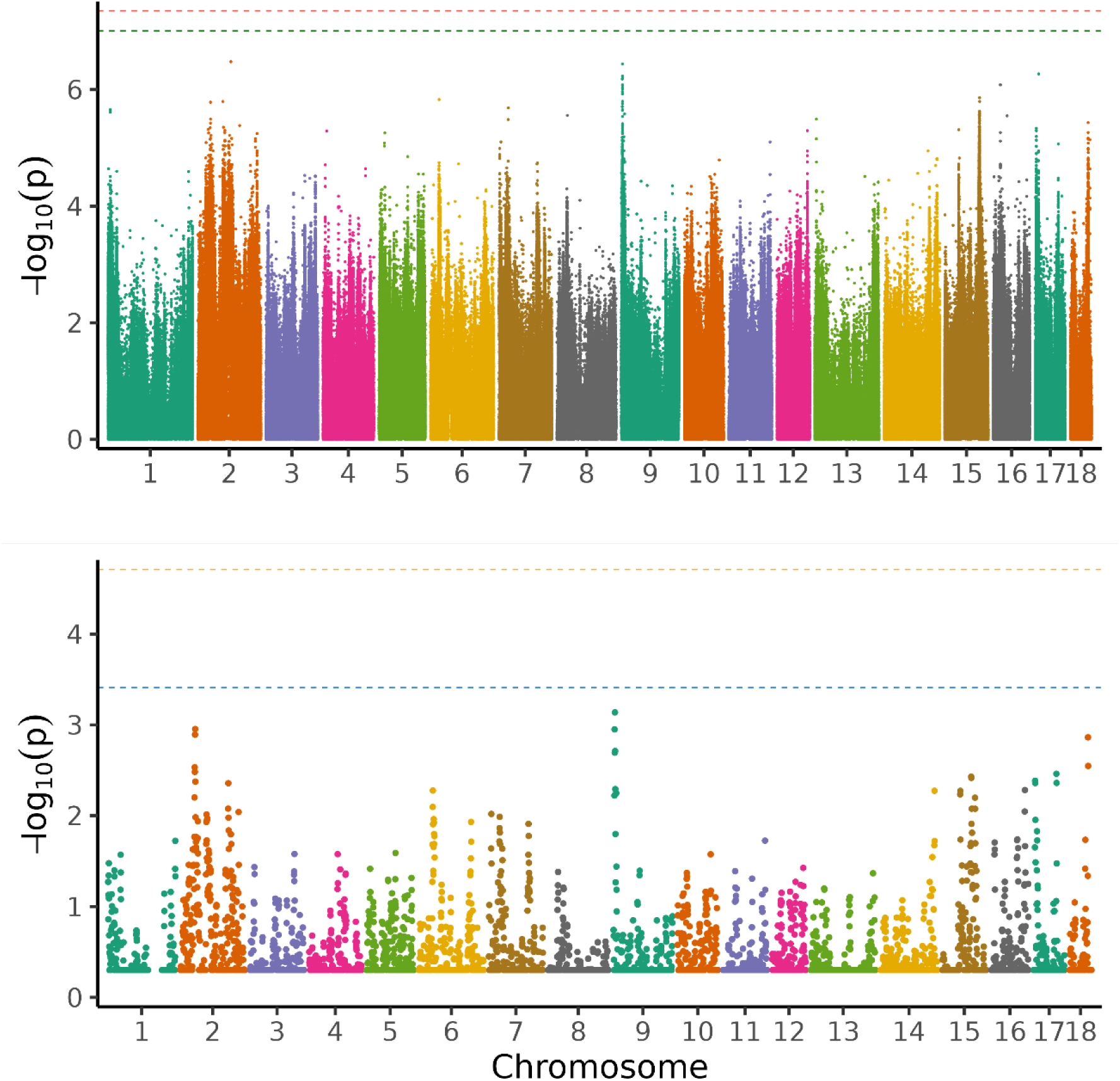
Manhattan plot of the genome-wide association analysis (above) and regional heritability mapping (below) of protein efficiency. The *x*-axis and the *y*-axis represent the chromosomes and the observed -log_10_(*P*-value), respectively. The red line is the threshold obtained by permutation, the green line in the Manhattan plot is the Bonferroni-corrected significance threshold at an alpha level of 0.05 (based on the number of sufficiently independent loci). In the lower panel, the orange line is the Bonferroni-corrected significance threshold at an alpha level of 0.05 (based on the number of unique windows tested) and the blue line is the suggestive threshold.

**Figure 2.**
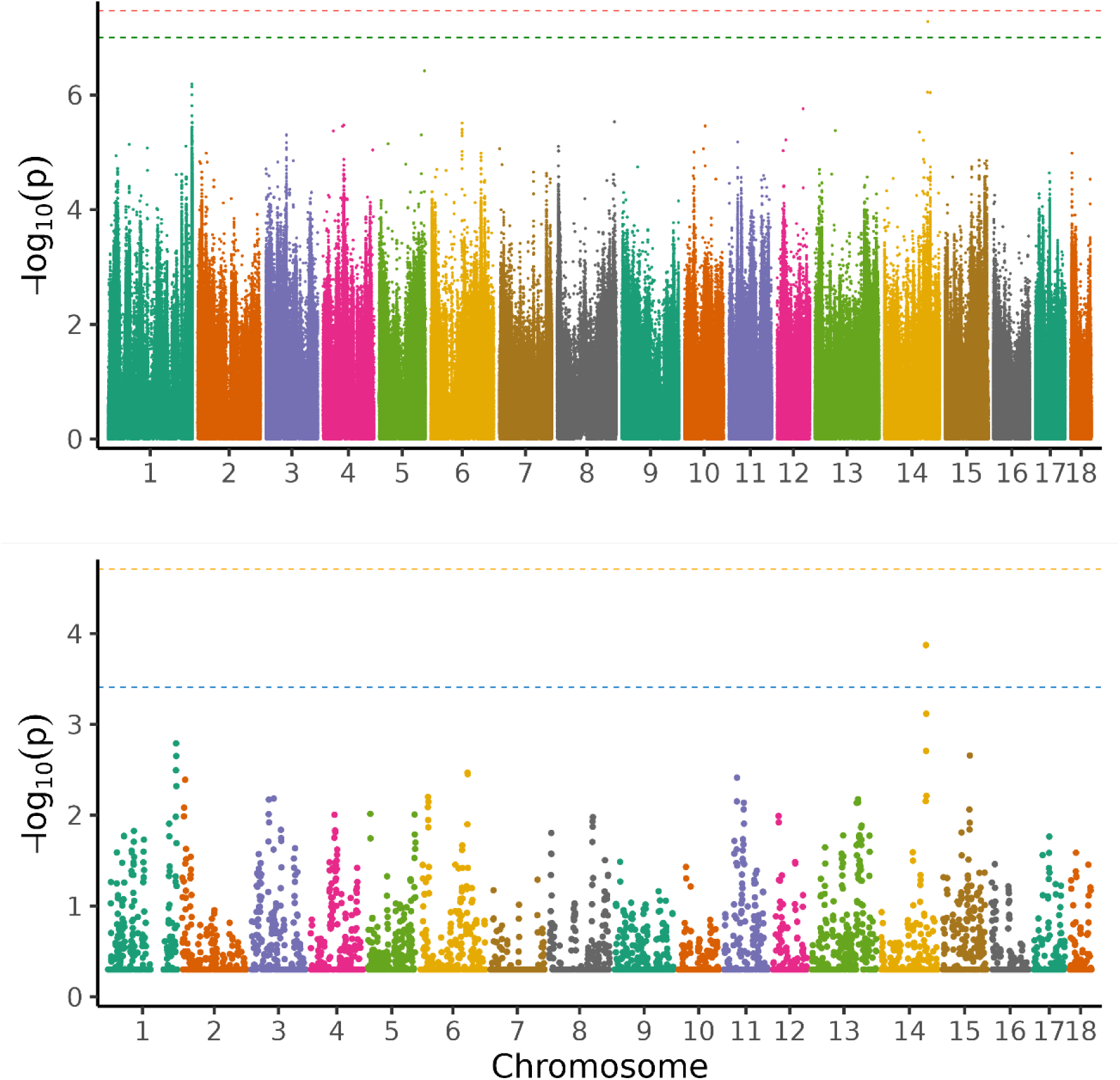
Manhattan plot of the genome-wide association analysis (above) and regional heritability mapping (below) of feed conversion ratio. The *x*-axis and the *y*-axis represent the chromosomes and the observed -log_10_(*P*-value), respectively. The red line is the threshold obtained by permutation, the green line in the Manhattan plot is the Bonferroni-corrected significance threshold at an alpha level of 0.05 (based on the number of sufficiently independent loci). In the lower panel, the orange line is the Bonferroni-corrected significance threshold at an alpha level of 0.05 (based on the number of unique windows tested) and the blue line is the suggestive threshold.

**Figure 3.**
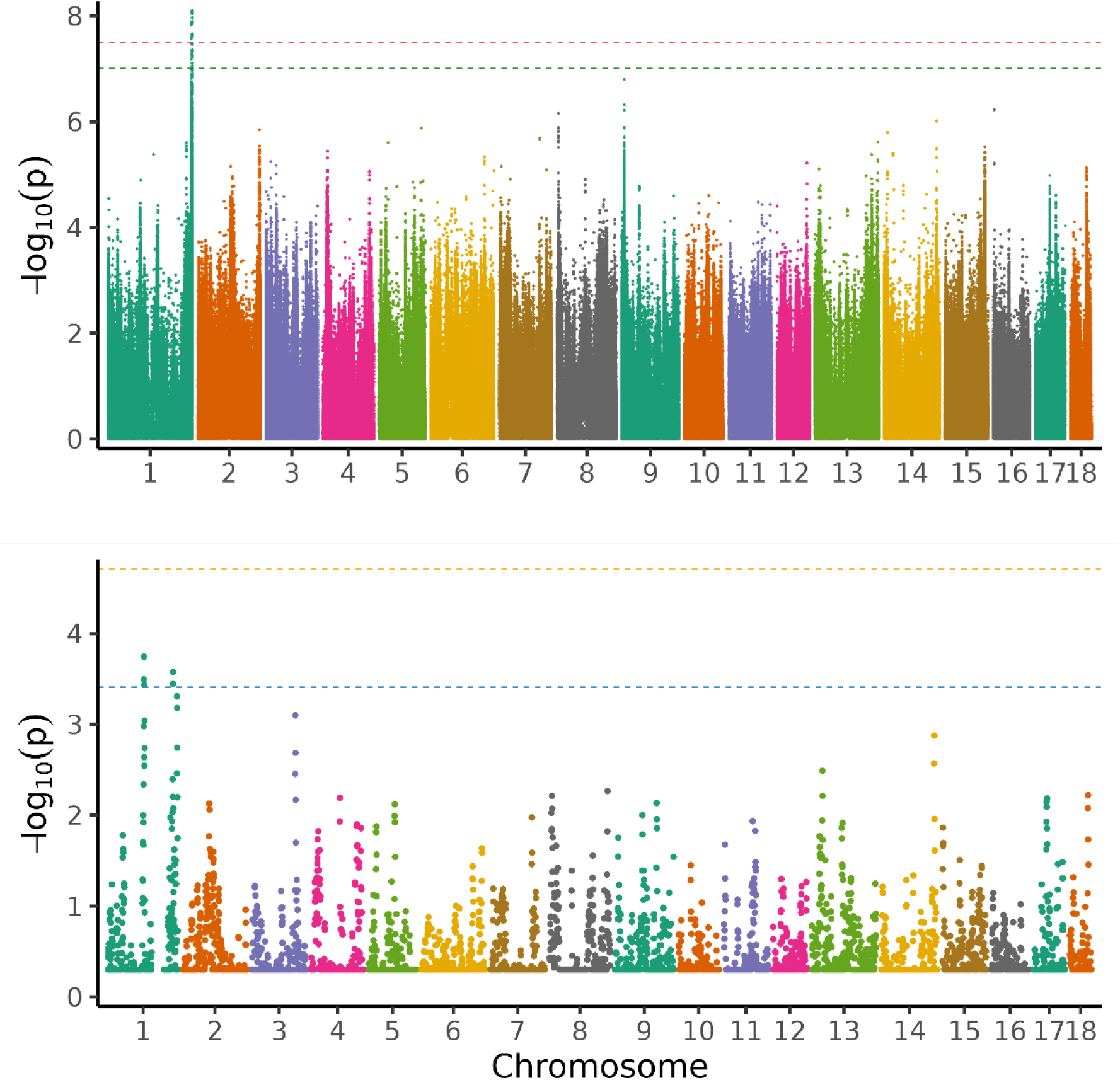
Manhattan plot of the genome-wide association analysis (above) and regional heritability mapping (below) of average daily gain. The *x*-axis and the *y*-axis represent the chromosomes and the observed -log_10_(*P*-value), respectively. The red line is the threshold obtained by permutation, the green line in the Manhattan plot is the Bonferroni-corrected significance threshold at an alpha level of 0.05 (based on the number of sufficiently independent loci). In the lower panel, the orange line is the Bonferroni-corrected significance threshold at an alpha level of 0.05 (based on the number of unique windows tested) and the blue line is the suggestive threshold.

**Figure 4.**
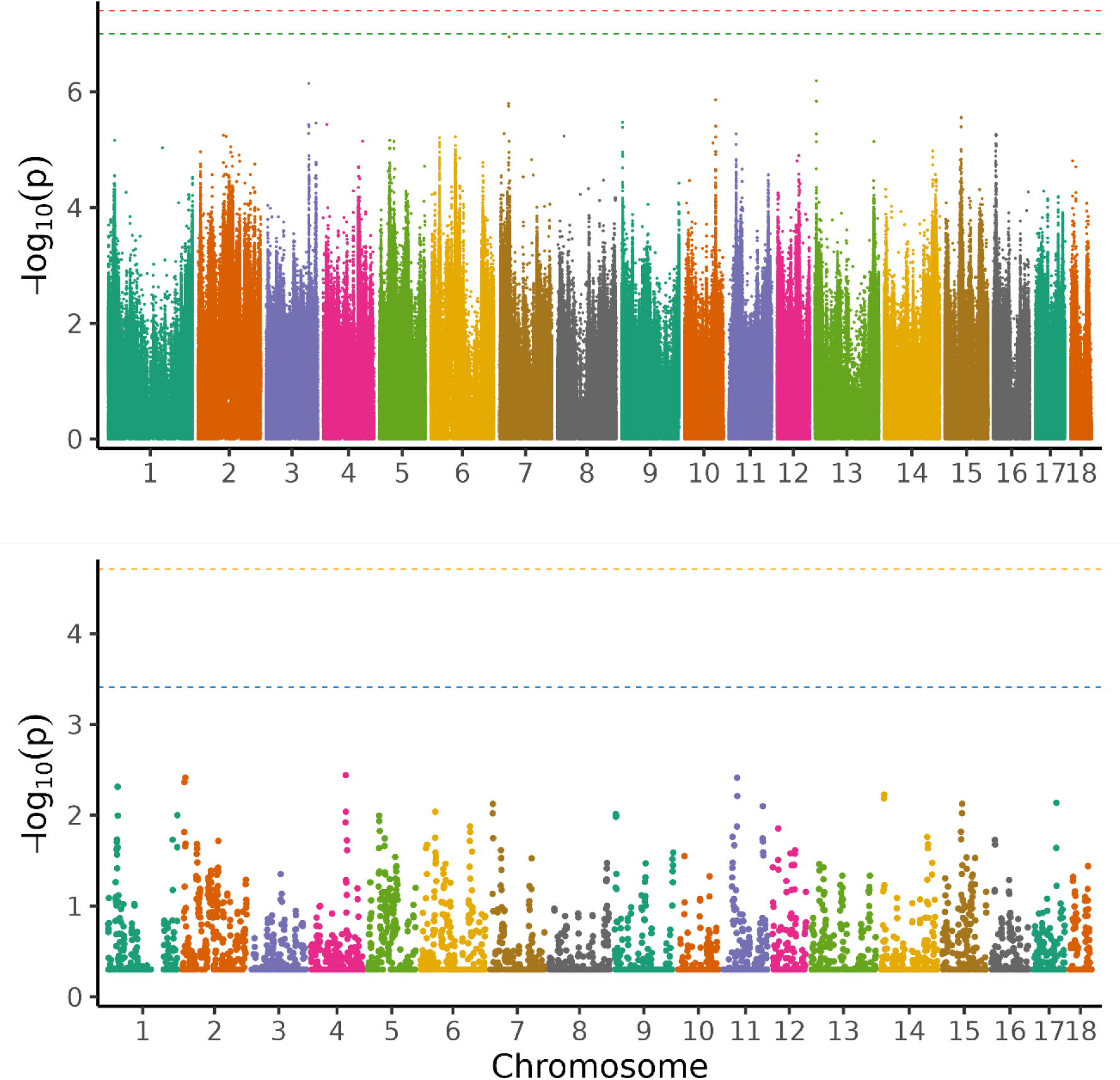
Manhattan plot of the genome-wide association analysis (above) and regional heritability mapping (below) of average daily feed intake. The *x*-axis and the *y*-axis represent the chromosomes and the observed -log_10_(*P*-value), respectively. The red line is the threshold obtained by permutation, the green line in the Manhattan plot is the Bonferroni-corrected significance threshold at an alpha level of 0.05 (based on the number of sufficiently independent loci). In the lower panel, the orange line is the Bonferroni-corrected significance threshold at an alpha level of 0.05 (based on the number of unique windows tested) and the blue line is the suggestive threshold.

**Figure 5.**
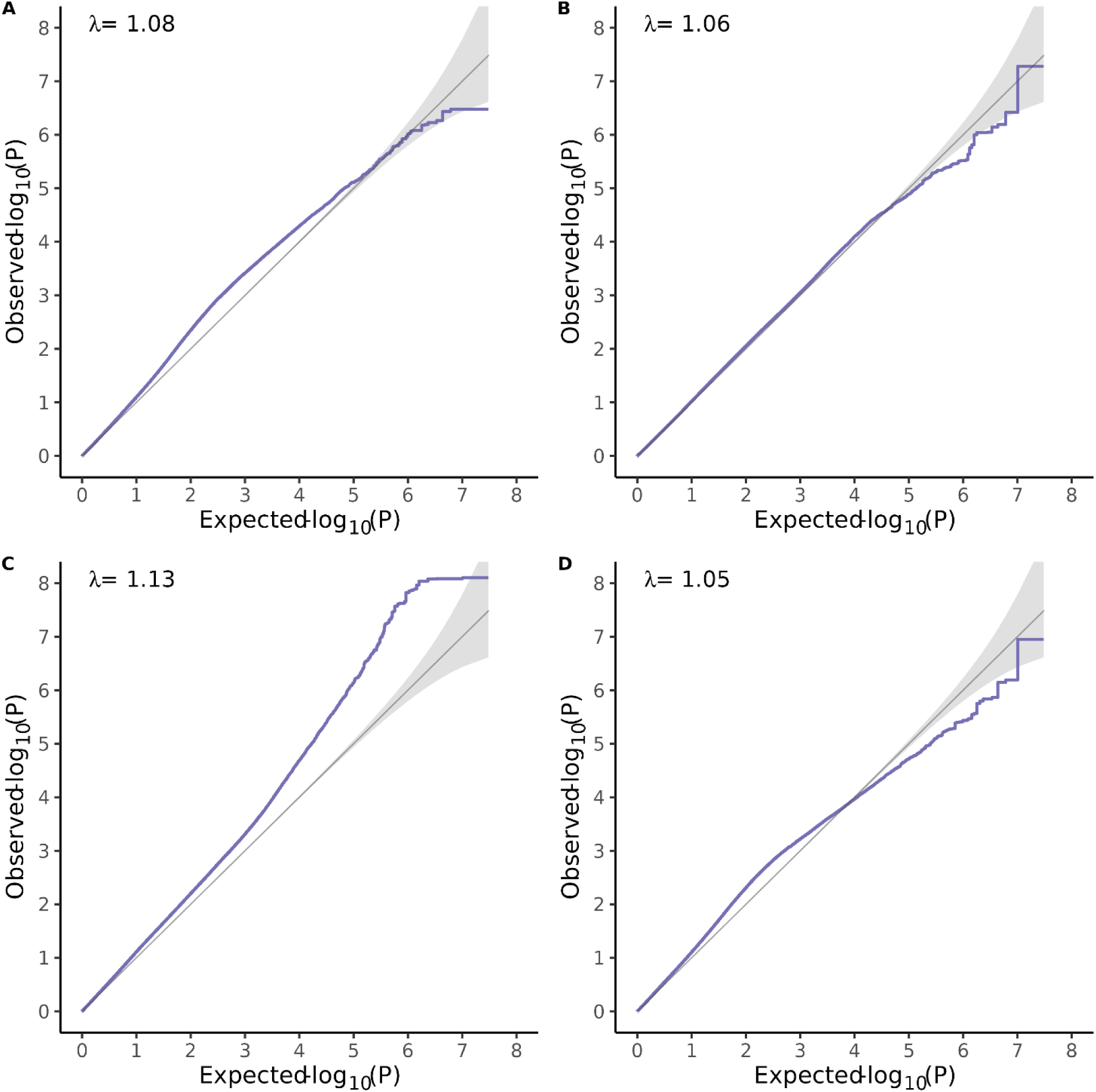
Quantile-Quantile plots. (A) protein efficiency, (B) Average daily gain (ADG), (C) Average daily feed intake (ADFI) and, (D) feed conversion ratio (FCR). λ is the genomic inflation factor.

**Table 1.**
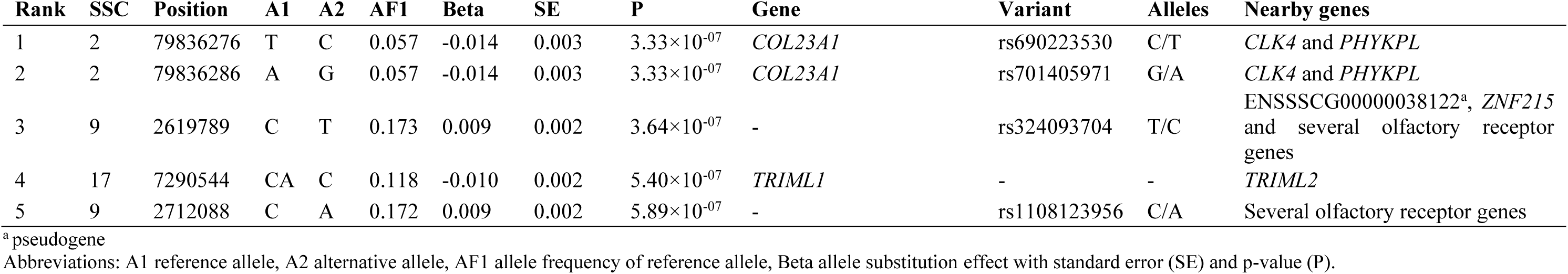
Top 5 variants from GWAS for protein efficiency with their position on the chromosome in base pairs, the gene they are located in, the variant ID and the alleles. Nearby genes (within a range of 100 kb up- and downstream) are also listed. Variants did not reach the permutation (4.47 × 10-8) nor the LD-pruned Bonferroni threshold (9.90 × 10-8).

| Rank | SSC | Position | A1 | A2 | AF1 | Beta | SE | P | Gene | Variant | Alleles | Nearby genes |
| --- | --- | --- | --- | --- | --- | --- | --- | --- | --- | --- | --- | --- |
| 1 | 2 | 79836276 | T | C | 0.057 | -0.014 | 0.003 | $3.33 \times 10^{-07}$ | *COL23A1* | rs690223530 | C/T | *CLK4* and *PHYKPL* |
| 2 | 2 | 79836286 | A | G | 0.057 | -0.014 | 0.003 | $3.33 \times 10^{-07}$ | *COL23A1* | rs701405971 | G/A | *CLK4* and *PHYKPL* |
| 3 | 9 | 2619789 | C | T | 0.173 | 0.009 | 0.002 | $3.64 \times 10^{-07}$ | - | rs324093704 | T/C | ENSSSCG00000038122<sup>a</sup>, *ZNF215* and several olfactory receptor genes |
| 4 | 17 | 7290544 | CA | C | 0.118 | -0.010 | 0.002 | $5.40 \times 10^{-07}$ | *TRIML1* | - | - | *TRIML2* |
| 5 | 9 | 2712088 | C | A | 0.172 | 0.009 | 0.002 | $5.89 \times 10^{-07}$ | - | rs1108123956 | C/A | Several olfactory receptor genes |

<sup>a</sup> pseudogene
Abbreviations: A1 reference allele, A2 alternative allele, AF1 allele frequency of reference allele, Beta allele substitution effect with standard error (SE) and p-value (P).

**Table 2.**
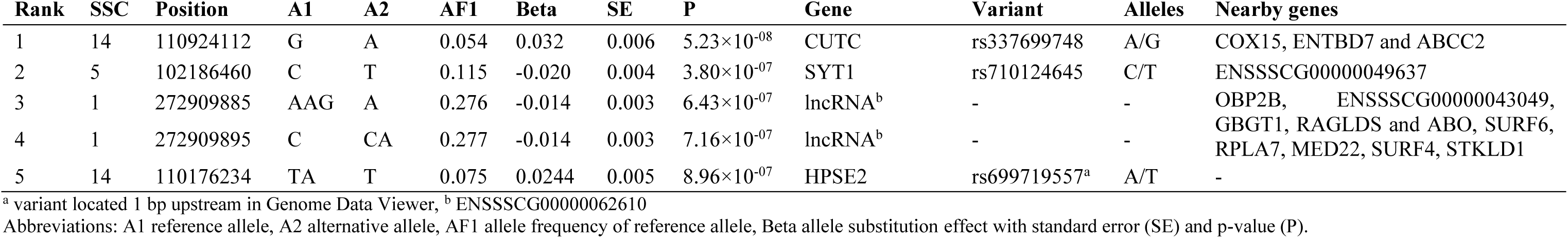
Top 5 variants from GWAS for average daily gain with their position on the chromosome in base pairs, the gene they are located in, the variant ID and the alleles. Nearby genes (within a range of 100 kb up- and downstream) are also listed. Variants ranked 2-5 did not reach the genome-wide (3.56 × 10^−8^) nor the suggestive threshold (9.93 × 10^−8^), but the first-ranking variant reached the suggestive threshold.

**Table 3.** Significant (in bold) and suggestive (in italics) variants from GWAS for average daily feed intake with their position on SSC1 in base pairs, the gene they are located in, the variant ID and the alleles. Nearby genes (within a range of 100 kb up- and downstream) are also listed. Variants ranked 1-26 reached the permutation threshold (3.19 × 10^−8^), and those ranked 27-45 the LD-pruned Bonferroni threshold (9.94 × 10^−8^).

| Rank | Position | A1 | A2 | AF1 | Beta | SE | P | Gene | Variant | Alleles | Nearby genes |
| --- | --- | --- | --- | --- | --- | --- | --- | --- | --- | --- | --- |
| 1 | 273496587 | T | G | 0.278 | 0.049 | 0.008 | $7.95 \times 10^{-09}$ | - | rs342474793 | G/T | VAV2 and BRD3 |
| 2 | 273496457 | A | G | 0.275 | 0.049 | 0.008 | $8.25 \times 10^{-09}$ | - | rs333113518 | G/A | VAV2 and BRD3 |
| 3 | 273496470 | T | G | 0.275 | 0.049 | 0.008 | $8.25 \times 10^{-09}$ | - | rs340291882 | G/T | VAV2 and BRD3 |
| 4 | 273496486 | C | T | 0.275 | 0.049 | 0.008 | $8.25 \times 10^{-09}$ | - | rs323875909 | T/C | VAV2 and BRD3 |
| 5 | 270982720 | C | A | 0.100 | 0.069 | 0.012 | $8.32 \times 10^{-09}$ | - | rs712027998 | A/C | FIBCD1 and LAMC3 |
| 6 | 273496625 | C | T | 0.278 | 0.049 | 0.008 | $8.34 \times 10^{-09}$ | - | rs326407596 | T/C | VAV2 and BRD3 |
| 7 | 273496743 | G | A | 0.275 | 0.048 | 0.008 | $9.16 \times 10^{-09}$ | - | rs325682162 | A/G | VAV2 and BRD3 |
| 8 | 273496749 | G | A | 0.275 | 0.048 | 0.008 | $9.16 \times 10^{-09}$ | - | rs339310280 | A/G | VAV2 and BRD3 |
| 9 | 273496759 | A | G | 0.275 | 0.048 | 0.008 | $9.16 \times 10^{-09}$ | - | rs320598057 | G/A | VAV2 and BRD3 |
| 10 | 273513647 | C | G | 0.248 | 0.051 | 0.009 | $1.08 \times 10^{-08}$ | BRD3 | rs321803809 | G/C | VAV2 and WDR5 |
| 11 | 273496721 | C | G | 0.275 | 0.048 | 0.008 | $1.29 \times 10^{-08}$ | - | rs344411671 | G/C | VAV2 and BRD3 |
| 12 | 270589733 | C | T | 0.172 | 0.055 | 0.010 | $1.33 \times 10^{-08}$ | - | rs322817388 | T/C | ASS1 |
| 13 | 273473392 | T | C | 0.247 | 0.050 | 0.009 | $1.38 \times 10^{-08}$ | VAV2 | rs333938622 | C/T | SARDH and BRD3OS |
| 14 | 270599545 | G | A | 0.161 | 0.056 | 0.010 | $1.39 \times 10^{-08}$ | - | rs326035908 | A/G | ASS1 |
| 15 | 270589693 | A | G | 0.172 | 0.055 | 0.010 | $1.50 \times 10^{-08}$ | - | rs335988374 | G/A | ASS1 |
| 16 | 270589695 | T | G | 0.172 | 0.055 | 0.010 | $1.50 \times 10^{-08}$ | - | rs340768103 | G/T | ASS1 |
| 17 | 273481971 | A | G | 0.245 | 0.050 | 0.009 | $2.20 \times 10^{-08}$ | - | rs323301985 | G/A | VAV2 |
| 18 | 273473590 | C | T | 0.243 | 0.049 | 0.009 | $2.38 \times 10^{-08}$ | VAV2 | rs335502076 | T/C | SARDH and BRD3OS |
| 19 | 273473601 | C | T | 0.243 | 0.049 | 0.009 | $2.38 \times 10^{-08}$ | VAV2 | rs319552596 | T/C | SARDH and BRD3OS |
| 20 | 273473603 | A | C | 0.243 | 0.049 | 0.009 | $2.38 \times 10^{-08}$ | VAV2 | rs713897779 | C/A | SARDH and BRD3OS |
| 21 | 273473613 | C | CG | 0.243 | 0.049 | 0.009 | $2.38 \times 10^{-08}$ | VAV2 | rs793814574 | CG/C | SARDH and BRD3OS |
| 22 | 273487853 | T | C | 0.250 | 0.049 | 0.009 | $2.43 \times 10^{-08}$ | - | rs335311042 | C/T | VAV2 and BRD3 |
| 23 | 272839507 | C | T | 0.472 | -0.041 | 0.007 | $2.47 \times 10^{-08}$ | LOC110256503<sup>a</sup> | rs3473060690 | T/C | GBGT1 |
| 24 | 270599467 | T | C | 0.161 | 0.055 | 0.010 | $2.69 \times 10^{-08}$ | - | rs699921403 | C/T | ASS1 |
| 25 | 270599489 | A | G | 0.161 | 0.055 | 0.010 | $2.69 \times 10^{-08}$ | - | rs713018859 | G/A | ASS1 |
| 26 | 270599528 | A | G | 0.161 | 0.055 | 0.010 | 2.69×10<sup>-08</sup> | - | rs706359608 | G/A | ASS1 |
| 27 | 273517030 | T | C | 0.249 | 0.049 | 0.009 | 3.32×10<sup>-08</sup> | BRD3 | rs333674056 | C/T | VAV2 and WDR5 |
| 28 | 270599666 | T | TATATAC | 0.162 | 0.055 | 0.010 | 3.43×10<sup>-08</sup> | - | rs713961016 | TATATAC/T | ASS1 |
| 29 | 273481384 | A | G | 0.248 | 0.049 | 0.009 | 3.47×10<sup>-08</sup> | - | rs318478034 | G/A | VAV2 and BRD3 |
| 30 | 273445392 | T | C | 0.250 | 0.049 | 0.009 | 4.29×10<sup>-08</sup> | VAV2 | rs325919278 | C/T | BRD3 |
| 31 | 273496990 | A | G | 0.273 | 0.047 | 0.009 | 4.65×10<sup>-08</sup> | - | rs340187248 | G/A | VAV2 and BRD3 |
| 32 | 273484075 | C | CAGGGA | 0.247 | 0.048 | 0.009 | 4.70×10<sup>-08</sup> | - | rs786847961 | CAGGGA/C | VAV2 and BRD3 |
| 33 | 273469880 | T | C | 0.249 | 0.049 | 0.009 | 4.85×10<sup>-08</sup> | VAV2 | rs328126130 | C/T | BRD3 |
| 34 | 270982734 | G | A | 0.101 | 0.065 | 0.012 | 5.34×10<sup>-08</sup> | - | rs1109265720 | A/G | FIBCD1 and LAMC3 |
| 35 | 273474175 | AG | A | 0.244 | 0.048 | 0.009 | 5.49×10<sup>-08</sup> | VAV2 | rs1107705322 | A/G | BRD3 |
| 36 | 273545902 | T | C | 0.242 | 0.048 | 0.009 | 5.54×10<sup>-08</sup> | - | rs320931279 | C/T | VAV2 and WDR5 |
| 37 | 273445528 | T | C | 0.251 | 0.048 | 0.009 | 5.70×10<sup>-08</sup> | VAV2 | rs329357351 | T/C | BRD3 |
| 38 | 273445540 | C | T | 0.251 | 0.048 | 0.009 | 5.70×10<sup>-08</sup> | VAV2 | rs338673132 | C/T | BRD3 |
| 39 | 273496268 | AATAG | A | 0.270 | 0.046 | 0.009 | 5.76×10<sup>-08</sup> | - | rs706632263 | A/AATAG | VAV2 and BRD3 |
| 40 | 273519510 | A | G | 0.249 | 0.048 | 0.009 | 6.02×10<sup>-08</sup> | BRD3 | rs691215421 | G/A | VAV2 and WDR5 |
| 41 | 270589753 | T | C | 0.171 | 0.053 | 0.010 | 6.67×10<sup>-08</sup> | - | rs332410580 | C/T | ASS1 |
| 42 | 273473786 | A | G | 0.242 | 0.048 | 0.009 | 7.87×10<sup>-08</sup> | VAV2 | rs344778327 | G/A | BRD3 |
| 43 | 273470949 | C | T | 0.248 | 0.048 | 0.009 | 8.76×10<sup>-08</sup> | VAV2 | rs324192941 | T/C | BRD3 |
| 44 | 273484432 | G | C | 0.251 | 0.047 | 0.009 | 8.79×10<sup>-08</sup> | - | rs329390136 | C/G | VAV2 and BRD3 |
| 45 | 273473274 | A | G | 0.246 | 0.047 | 0.009 | 9.74×10<sup>-08</sup> | VAV2 | rs325075572 | G/A | BRD3 |

<sup>a</sup> non-coding RNA
Abbreviations: A1 reference allele, A2 alternative allele, AF1 allele frequency of reference allele, Beta allele substitution effect with standard error (SE) and p-value (P).

**Table 4.**
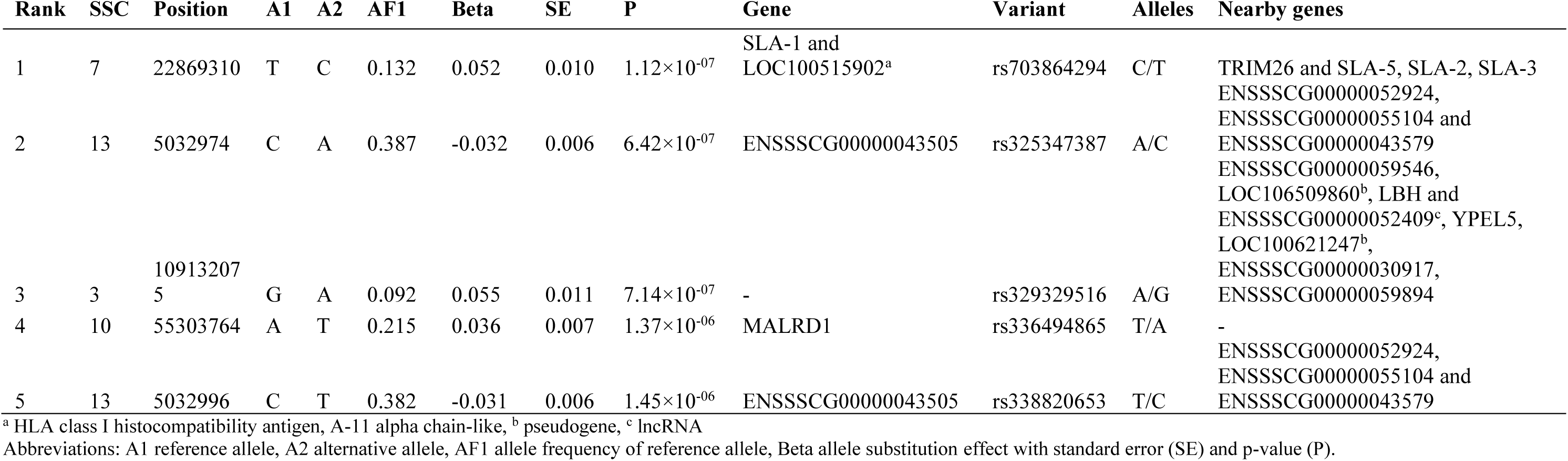
Top 5 variants from GWAS for feed conversion ratio with their position on the chromosome in base pairs, the gene they are located in, the variant ID and the alleles. Nearby genes (within a range of 100 kb up- and downstream) are also listed. None of the variants reached the permutation (3.93 × 10^−8^) or the LD-pruned Bonferroni threshold (9.91 × 10^−8^).

**Table 5.** Descriptive statistics and pedigree and genomic heritability estimates for protein efficiency and performance traits.

| Trait | N | Mean ± SD | Min - Max | $h^2_{ped}$ (SE) | $h^2_{geno}$ (SE) |
| --- | --- | --- | --- | --- | --- |
| PE | 1025 | 0.39 ± 0.03 | 0.28 – 0.49 | 0.54 (0.10) | 0.42 (0.05) |
| ADG | 1033 | 0.85 ± 0.11 | 0.51 – 1.20 | 0.45 (0.11) | 0.47 (0.04) |
| ADFI | 1034 | 2.26 ± 0.31 | 1.30 – 3.14 | 0.53 (0.12) | 0.42 (0.05) |
| FCR | 1024 | 2.68 ± 0.21 | 1.99 – 3.77 | 0.39 (0.12) | 0.33 (0.04) |

PE: Protein efficiency; ADG: average daily gain; ADFI: average daily feed intake; FCR: feed conversion ratio.

### Regional heritability mapping

In total, around 5,230 genomic regions were subjected to RHM for each trait. For all traits, no region was found significant at the genome-wide level (Figs. 1 to 4). No region reached the suggestive threshold for PE (Fig. 1, Table S5). There was one region on SSC14 that passed the suggestive threshold for ADG with a heritability of 0.15 (Fig. 2, Table S6). Five regions reached the suggestive threshold for ADFI on SSC1 (Fig. 3, Table S7). These were three overlapping windows (635000_640000, 637500_642500 and 640000_645000), as well as two overlapping windows (1155000_1160000 and 1157500_1162500). The heritabilities in these five windows were rather low (0.06 to 0.1). For FCR, there were no regions that passed either threshold (Fig. 4, Table S8) and heritabilities of windows were low throughout (0.008 to 0.04).

### Support of GWAS variants by RHM

For PE, the two methods seemed to agree quite well and confirmed each other (Fig. 1). The closest variants to the LD-corrected Bonferroni threshold (-log_10_(P) = 7.00) in GWAS were on SSC2, SSC9 and SSC17. Also in RHM, the windows that were closest to the suggestive threshold (-log_10_(P) = 3.41) were on SSC2 and SSC9. However, the hightest ranking variants in GWAS on SSC2 were not included in the highest ranking windows of RHM, but three other variants in the GWAS top 50 (ranked 26-28) were included in window 205000_210000, which was ranked 10 in RHM (Tables S1 and S5). This window overlapped with two others on SSC2 in the top ten windows (205000_210000, 207500_212500 and 210000_215000; 10’000 variants), spanning the region of 30,144,793 bp to 32,170,603 bp. The highest-ranking variant on SSC9 was also not found in a RHM window in the top 50, but 17 variants on SSC9 (ranking 5 to 33) were present in the top-ranking window in RHM (20000_25000; Table S5). Two adjacent windows were top ranking on SSC9 (15000_20000 and 20000_25000), which encompassed 10,000 variants and spanned the region 2,013,927 bp to 2,730,410 bp. The suggestive variant on SSC14 in GWAS for ADG was not present in a top 50 window in RHM and the suggestive window on SSC14 (637500_642500) did not contain any variant in the top 50 (Fig. 2; Table S2 and S6). However, three other variants among the top 10 in GWAS on SSC14 were found in the adjacent window ranking 18^th^ (635000_640000; both windows ranging from 117,072,833 bp to 118,516,125 bp). All 19 variants on SSC1 in the top 50 were present in two overlapping windows in RHM (1227500_1232500 and 1230000_1235000; from 272,754,902 bp to 273,355,069 bp, spanning 7,500 variants). Two variants on SSC1 in the top 5 were found in 2 overlapping windows among the top 5 (Table 2, Table S6). The variant on SSC5 ranked 2^nd^ in GWAS was present in the 30^th^ ranking window in RHM (662500_667500). All 45 variants that passed one of the thresholds in GWAS for ADFI belonged to two overlapping windows in RHM (1232500_1237500 and 1235000_1240000) on SSC1 within the region of 273,495,531 bp to 273,690,809 bp (Fig. 3, Table S3 and S6). In general, the pattern for FCR was more inconsistent than for the other traits, with no overlap of top 5 variants and windows (Fig. 4, Table S4 and S8). For instance, one variant on SSC7 was rather close to the suggestive threshold (-log_10_(P) = 7.00) in GWAS, but it was not present in any window in the RHM top 50.

### Genes related to genomic variants and Regional Heritability Mapping windows

The first two variants from GWAS for PE were located on SSC2at 79,836,276 bp and 79,836,286 bp (Table 1). These variants were found within the *COL23A1* gene, and were approximately 13-36 kb downstream from the start and end of *CLK4* and 977 bp upstream from *PHYKPL*. The third and the fifth ranking variants were located on SSC9 at 2,619,789 bp and 2,619,789 bp, respectively. Both variants were within an intergenic region and flanked by four different olfactory receptor genes. The fourth lead variant was on SSC17 in the *TRIML1* gene. Genes located in the leading windows in RHM for PE are shown in Table S9. Several protein coding genes, lncRNAs and one pseudogene were located in the two adjacent windows on SSC9, such as *PPFIBP2, OLFML1*, *SYT9, RBMXL2, NLRP14, ZNF215,* and 16 olfactory receptor genes. The three overlapping windows on SSC2 contained protein coding genes, such as *MPPED2*, *ARL14EP*, *FSHB*, *KCNA4* and *METTL15*, as well as 25 lncRNAs and 3 pseudogenes. For ADG, the suggestive region identified by GWAS on SSC14 at 110,924,112 bp was situated in the *CUTC* gene (Table 2). The second-ranked variant on SSC5 at 102,186,460 bp was in the *SYT1* gene. Variants ranked 3 and 4 were found within an intergenic region on SSC1 between *OBP2B* and *ABO*. The *HPSE2* gene on SSC14 harboured the fifth-ranking variant (110,176,234 bp). The suggestive variant was not found in a window reaching the significance or suggestive threshold in RHM, but the suggestive window on SSC14 contained the *SORCS1* gene, 2 noncoding RNAs, 6 lncRNAs and one small nucleolar RNA (Table S10). Three further variants (ranks 6-8) in GWAS (Table S2) were located within the *SORCS1* gene, all within one RHM window (635000_640000) spanning 117,072,833 to 118,069,574 bp on SSC14. Several variants on SSC1 were included in overlapping windows (1227500_1232500 and 1230000_1235000), which contained 25 protein coding genes, 1 noncoding RNA, 7 lncRNAs and 5 small nucleolar RNA (Table S10).

Three variants for ADFI on SSC1 (one significant and two suggestive) were located in the *BRD3* gene, five significant (located between 273.4733 Mb – 273.4736 Mb) and eight suggestive variants in the *VAV2* gene, and one in the uncharacterized LOC110256503, a non-coding RNA, which is 11.58 to 8.516 kb downstream of the *GBGT1* gene (Table 3). The remaining 19 significant and 9 suggestive variants were located in the vicinity of *VAV2* and *BRD3*, *FIBCD1* and *LAMC3*, *ASS1*, *SARDH* and *BRD3OS* (*BRD3* opposite strand), and *WDR5*. *VAV2* was approximately 8 kb downstream from *SARDH* and 84 kb upstream from *WDR5*. The two RHM windows from 273,498,257 to 273,634,463 bp containing all significant and suggestive variants spanned also *BRD3*, *BRD3OS*, *WDR5*, as well as six lncRNA, a snRNA (RNU6ATAC), one miRNA (MIR9812) and another noncoding RNA (Table S11). The genes in the three overlapping RHM windows ranging from 103,336,199 to 105,376,226 bp and two overlapping windows ranging from 260,888,411 to 262,071,057 bp on SSC1 passing the suggestive threshold harboured 30 protein coding genes (for example *MBD2*, *STARD6*, *TCF4*, *PSMD5* and *RAB14*), six pseudogenes, 15 lncRNAs and one snRNA (Table S11). The close-to-suggestive variant for FCR on SSC7 was located in the SLA-1 gene and in a HLA class I histocompatibility antigen, A-11 alpha chain-like gene (LOC100515902), near TRIM26 and SLA-5, SLA-2, SLA-3, coding for several Scr-like adapter proteins. Variants ranked 2 and 5 on SSC13 were located in ENSSSCG00000043505, with no described genes nearby, and the third and fourth-ranking variants were located in an intergenic region and in the MALRD1 gene, respectively.

## Discussion

Here, we aimed to identify genomic regions associated with PE, ADG, ADFI and FCR using both genome-wide association studies (GWAS) and regional heritability mapping (RHM). We observed differences in the strength of evidence supporting associations between genomic variants or regions and phenotypes across different traits. While we identified several variants and windows at both the significant and suggestive thresholds for ADFI and at the suggestive threshold for ADG, the evidence for variants and windows underlying PE, although present below the threshold in both GWAS and RHM individually, increased our confidence when the results were integrated. In contrast, this combined approach failed to uncover any clear genomic regions associated with FCR.

### Genes associated with traits

#### Average daily feed intake

We identified 26 significant variants for ADFI, and a further 19 variants that reached the suggestive threshold, all on SSC1 within the region of 270-273.5 Mb. Our study also found five suggestive regions associated with ADFI located at 103-105 Mb and 260-262 Mb on SSC1. Nosková et al. [31] also detected a QTL associated with ADFI on SSC1 at the 270–272 Mb position in the same population. This chromosomal region also harbours QTL for ADFI in other pig breeds and populations [63, 64, 65], suggesting an ancestral origin of this QTL.

Most variants within a gene mapped to *VAV2*, which plays a key role in immunity by preventing bacterial attachment and subsequent uptake into cells [66]. The gene is also involved in pathways critical for skeletal muscle growth and metabolic regulation via the IGF1-insulin pathway. VAV2 deficiency in mice led to reduced muscle mass, insulin responsiveness and ultimately symptoms resembling metabolic syndrome [67]. In pigs, *VAV2* expression in muscle and liver responded to different dietary fatty acid profiles [68]. The *BRD3* gene also contained several ADFI-associated variants. It encodes a protein that influences the expression of other genes by increasing their accessibility to RNA polymerase II for transcription [69], and regulates the expression of several immunity-related genes, thereby constituting an important part of the innate immunity [70]. In pigs, *BRD3* has been shown to be involved in the immune response to African swine fever [71] and porcine endogenous retrovirus [72]. The level of methylation of *BRD3* in the jejunum of piglets has been suggested to play a role in gut maturation and adaptation to bacterial colonisation associated with milk feeding [73].

Intergenic variants were located near *FIBCD1, LAMC3* and the lncRNA ENSSSCG00000042995. *FIBCD1* encodes a chitin-binding receptor with functions in the innate immune system. It responds to helminth antigens, helping intestinal cells control fungal colonization and shape the composition of the gut microbiome [74]. This role is further supported by findings from a mouse model, where overexpression of *FIBCD1* mitigated chemotherapy-induced weight loss, probably by reducing mucositis and gastrointestinal dysbiosis [75]. In pigs, a SNP close to *FIBCD1* was associated with the lean meat content of ham [76]. *LAMC3*, part of the extracellular matrix pathway (GO: 0031012), influences tissue development, repair, differentiation and cell migration [77]. As such, it is implicated in adipose tissue remodelling, inflammation and insulin resistance associated with obesity [76]. In pigs, its expression in adipose tissue was associated with fatness traits [78] and varied in muscle depending on dietary fat content [79].

*ASS1*, another gene in the vicinity of several variants in an intergenic region, encodes the urea cycle enzyme argininosuccinate synthetase, which is required to convert neurotoxic ammonia to urea in the liver. Deficiency of this gene is associated with metabolic disorders such as citrullinemia [80], insulin resistance and diabetes [81] in humans. It contributes to the innate immune responses to viral [81] and bacterial infections, with expression changes observed in a porcine liver injury model after LPS challenge [82]. Due to its role in the response to growth hormone (GO:0060416) process, *ASS1* is a likely candidate gene for growth and body size in pigs [83, 84]. It has also been proposed as a candidate gene for feed efficiency in pigs, probably as a host genetic effect [84]. *ASS1* expression in newborn piglets responded dynamically to antibiotics and maternal fecal microbiome transplant interventions, influencing alanine biosynthesis and amino acid metabolism [85]. Through the biological process’response to nutrient’ (GO:0007584), this gene may directly regulate feed intake by sensing the nutrient composition of the diet and adapting intake to the body’s nutrient requirements. This is further supported by the lower expression observed in the umbilical vein of intrauterine growth-retarded piglets, likely due to reduced fetal nutrient supply [86]. It is also linked to energy production and fat storage via the cellular response to oleic acid (GO:0071400). Higher *ASS1* expression was observed in the semimembranosus muscle of fatter Pulawska pigs compared to the leaner Polish Landrace [87], and its expression was positively correlated with fat area between the 13th and 14th rib of the longissiumus dorsi muscle [88].

We identified a significant variant in the uncharacterised lncRNA LOC110256503 (located between 272,836,566 and 272,843,572 on SSC1), which is partially overlapping ENSSSCG00065003108.1 (272,839,760 - 272,843,574). It is located near *GBGT1*, which starts around 8 kb upstream and encodes the Forssman synthase, a member of the ABO gene family, which determines blood group antigens through glycosyltransferase activity. This gene is likely inactive in pigs, as in humans and other species [89, 90] suggesting it is unlikely to be the regulatory target of this lncRNA. Given that lncRNAs can regulate the transcription, translation and metabolism of a wide range of genes and their products, including those distant from their own genomic location [91], detailed investigation is required to identify the regulatory targets of this lncRNA and to determine whether its function is linked to genes associated with traits such as ADFI.

Another intergenic variant for ADFI was located near *WDR5*, 75 kb downstream of the start of *BRD3*. It regulates gene expression through histone modification and has a broad range of functions in development, cell division, signal transduction, vesicular trafficking, cytoskeletal assembly, cell cycle control, and apoptosis [92, 93]. It is the target of a long intergenic noncoding RNA (MSTRG.2530) associated with differences in skeletal muscle growth between Yorkshire and Tibetan pigs [94]. An ADFI variant (position 270,982,734 on SSC1) is located 21 kb upstream of *LAMC3* and 65 kb downstream of the end of *QRFP*, encoding a hypothalamic neuropeptide that is highly evolutionarily conserved. It influences feeding behaviour and glucose homeostasis through G protein-coupled receptors [95, 96]. Administration of the *QRFP* peptide directly into the brain of mice resulted in increased feed intake and foraging behaviours [97]. It has been proposed as a prohormone gene (the precursor molecule of a neuropeptide) in pigs [98], and may underlie the reduced backfat thickness observed in Pietrain compared to other pig breeds [99]. A selection signature related to backfat thickness at the shoulder in Yorkshire pigs harboured this gene [100] and a variant associated with residual feed intake was located near it [101].

The three overlapping suggestive RHM windows (103,336,199-105,376,226 bp) on SSC1 contained protein-coding genes with roles in pig production traits, including *MBD22*, *STARD6*, *RAB27B* and *TCF4*. *MBD2* is involved in the differentiation of porcine mesenchymal stem cells into adipocytes [102] and regulates intramuscular fat content and fatty acid composition in the pig longissimus dorsi muscle [103]. *STARD6* binds cholesterol and other steroids and plays a role in lipid transport and metabolism [104, 105]. *RAB27B*, a target of the lncRNA MSTRG.28207.43, was significantly upregulated in the colon during heat stress-induced intestinal inflammation in pigs [106]. The transcription factor *TCF4* regulates myogenesis [107] and is involved in intestinal epithelium [108] and muscle development in pigs [109]. The two overlapping suggestive windows on SSC1 at 260,888,411 to 262,071,057 bp harboured genes with multiple documented roles in pigs, such as *PSMD5*, which was differentially expressed in the blood of piglets from lines differentially selected for RFI [110], and, *PHF19*, both of which were found in a selection signature in Indian pigs [111]. The gene *TRAF1*, involved in antiviral response and innate immunity [112], was associated with viral load in PRRS-infected pigs [113]. Another innate immunity-related gene, *C5*, was implicated in differences in monocyte levels in pig blood [114]. *RAB14*, which is involved in the regulation of hepatic insulin signalling and glucose metabolism [115], was also found in co-expression networks for skeletal muscle myogenesis across species [116].

#### Average daily gain

For ADG, we identified a suggestive variant in the *CUTC* gene on SSC14. It is an evolutionary conserved copper transporter protein, important for maintaining intracellular copper ion homeostasis [117]. In pigs, Sahana et al. [118] reported a FCR-associated SNP near the *CUTC* gene. In Nelore cattle, the gene harbors an allele-specific expression QTL in the longissimus thoraci muscle that might affect meat quality traits [119]. The suggestive region identified by RHM on SSC14 contained *SORCS1*, which is implicated in obesity-induced type 2 diabetes [120, 121]. It is a sorting receptor that helps direct proteins to their correct locations in cells, and is found in hypothalamic neurons, where it attenuates signaling by *BDNF* (brain-derived neurotrophic factor), thereby regulating energy homeostasis [122]. The knock-out of *SORCS1* together with *SORCS3* in mice leads to the overproduction of an orexigenic neuropeptide and results in increased food intake, decreased locomotor activity, reduced use of lipids as metabolic fuel and increased adiposity, despite overall reduced body weight [122]. *SORCS1* was suggested as candidate gene for backfat thickness in pigs [123], fat deposition in beef cattle across several breeds [124], and FCR in Pekin ducks [125].

#### Protein efficiency

Neither GWAS nor RHM yielded significant or suggestive results for PE, but both methods suggested the potential involvement of variants on SSC2 and SSC9. Previously, Shirali et al. [25] found associations on SSC2 and SSC9 for nitrogen excretion traits during the 60 – 140 kg growth stage, but it is unclear whether these associations were present in the same region as those for PE in our study as they used a different reference genome. The two top-ranking variants on SSC2 in GWAS were located in *COL23A1*, with the genes *CLK4* and *PHYKPL,* nearby (Table 2). *COL23A1* was associated with meat quality [126] and cholesterol levels [127] in pigs and is differentially methylated in humans with regard to waist-to-hip ratio [128] and body mass index [129]. In men, but not women, this gene is associated with weight gain after weight loss [130]. Both *PHYKPL* and *COL23A1* were implicated in nitrogen metabolism and nitrogen excretion in lactating Holstein-Friesian cows [131]. Considering the role of these two genes in nitrogen utilization, albeit in a different species, and in metabolism and growth, they may be considered as potential candidate genes in protein efficiency in pigs.

The three overlapping RHM windows on SSC2 (30,144,793 – 32,170,603 bp) contained 16 (long) noncoding RNAs and, amongst others, the protein-coding genes *MPPED2*, *ARL14EP*, *FSHB*, *KCNA4* and *METTL15*. In humans, *MPPED2* was associated with kidney function [132], and, in pigs, it was contained in a gene-co-expression module in muscle that differed between lean and obese breeds [133]. *KCNA4* has been reported in relation to production traits in pigs [134]. *METTL15* is required for mitochondrial protein synthesis [135] and linked to blood phosphorus levels in pigs [136]. Considering the reported functions of these genes, it is unclear whether they are plausible candidates for PE. It seems more likely that the noncoding RNAs in the highlighted region may a significant role in shaping the PE phenotype, although their targets remain to be identified.

The top-ranking variants for PE on SSC9 were located in intergenic regions (Table 2). We examined the combined region (2,013,927 to 2,812,088 bp) that included the overlapping top-ranked windows and extended 100 kb upstream and downstream of the variants (Table S11). In this region we identified several genes, including *PPFIBP2*, *OLFML1*, *SYT9*, *RBMXL2*, ENSSSCG00000060118 (largely overlapping with *RBMXL2*), *NLRP14*, LOC100524097, the transcription factor *ZNF215* and a set of olfactory receptor genes. Some of these genes were linked to feed efficiency, muscle and adipose tissue metabolism, body weight, milk protein, immune function and other relevant traits. *PPFIBP2*, involved in neuromuscular junction development (GO:0007528), was among the differentially expressed genes among Polish pig breeds [137] and upregulated in pale, soft, and exudative meat [138]. It was also differentially expressed in a non-alcoholic fatty liver pig model [139] and hypermethylated in the muscle of obese rabbits [140]. It has been proposed as a candidate gene for lactation persistency [141] and milk protein yield in buffalo [142], as well as milk protein percentage [143] and carcass traits [144] in cattle. *SYT9,* regulating dopamine secretion (GO:0014059) and exocytosis (GO:0070382 and GO:0017156, amongst others), shows maternal transmission ratio distortion in Entrepelado pigs, although the resulting phenotype remains unclear [145]. It was identified as candidate genes for shear force in pork [126], monounsaturated fatty acid content of the longissimus dorsi muscle in Ningxiang pigs [146] and lactation persistency in buffalo [141].

*RBMXL2* has been proposed as a candidate gene for postnatal growth [147] and milk protein [143] and is contained in a region identified as selection signature linked to domestication in cattle [148]. In buffalo, it is associated with lactation persistency [141]. *NLRP14*, involved in the regulation of inflammatory response (GO: 0050727), was associated with feed efficiency in cattle [149,150]. It is highly expressed in the porcine intestine, but its predicted transcript in pigs contains multiple stop codons, indicating it may be an expressed pseudogene [151]. *ZNF215*, is a well-known imprinted gene in vertebrates [88,89], has been identified as a candidate for shear force in pork [126] and milk protein in Holstein [152].

Olfactomedin proteins, such as *OLFML1*, are expressed throughout the brain and are key for the early development of the nervous system (neural crest), and are implicated in obesity [153] and non-alcoholic fatty liver disease [154, 155]. The region on SSC9 highlighted by both GWAS and RHM as potentially involved in PE contained 16 olfactory receptor genes. Olfactory transduction pathways and the olfactory receptor genes involved have been linked to residual feed intake in pigs [156], reflecting the extraordinary importance of olfaction in their foraging behaviour [157]. Olfactory pathways interact with satiety hormones like ghrelin, orexins, neuropeptide Y, insulin, leptin, and cholecystokinin [158]. In pigs, serum leptin correlated strongly with RFI [159], and plasma leptin was significantly higher in high efficiency lines [160].

#### Mechanisms underlying feed intake, growth and efficiency

The mechanisms by which the identified genes influence ADFI, ADG, and PE phenotypes range from direct effects, such as regulating feeding behavior (*QRFP*, *SORCS1*, *SYT9*, olfactory receptors) and nutrient sensing (*ASS1*), to indirect effects on immune function (*VAV2*, *BRD3*, *FIBCD1*, *ASS1*, *TRAF1*, *C5*), gut microbiome composition (*BRD3*, *FIBCD1*), and metabolism (*VAV2*, *LAMC3*, *ASS1*, *SORCS1*, *COL23A1*, *MPPED2*, *PPFIBP2*, *SYT9*, *OLFML1*, *MBD2*, *STARD6*, *RAB14*). Several genes were linked to the urea cycle (*ASS1*) and protein utilization in dairy cattle (*COL23A1*, *PHYKPL*, *PPFIBP2*, *RBMXL2*, *ZNF215*), highlighting their relevance for improving PE and sustainability in livestock. In PigGTEx, considerable expression of these genes in relevant tissues (e.g. muscle, adipose tissue, brain, gastrointestinal tract), as well as a large number of is molecular QTL and lncRNAs were documented. Apart from processes influencing gut microbiome composition and metabolism, pathways involved in nutrient sensing and regulation of feed intake may be of particular relevance to PE. Studies suggest that pigs, if given the choice between diets differing in protein content, actively select a diet with a protein content that appeared to optimize their growth [161, 162] but the generalisation of these results and the exact underlying mechanisms remain unresolved.

Many of the highlighted genes overlap with a QTL on SSC1 for production traits in Swiss Large White pigs [31], including *ASS1*, *FIBCD1*, *LAMC3*, *QRFP*, and the lncRNA ENSSSCG00000042995, supporting their role as candidate genes for ADFI and related traits. However, another QTL (157.7–162.7 Mb) identified by Noskova et al. [31] was not detected in our study, likely due to differences in sample size, phenotyping, and diet composition. Noskova et al. [31] used deregressed progeny phenotypes with a ∼5.5× larger sample size, while our study measured feed intake directly for all animals on an experimental farm. Importantly, the diets differed significantly: Noskova et al. [31] used a commercial diet optimized for performance, whereas the diet in the current study was formulated to mimic future sustainable feeding strategies by including more native protein feed crops. This diet also had a reduced crude protein and essential amino acid content (described in detail in [10]), and may have differed slightly in other aspects, such as digestible energy content (known to regulate feed intake in pigs [163]), fatty acid composition and fibre content. Interestingly, *VAV2*, highlighted in our study for its numerous variants, was absent from the analysis of Noskova et al. [31]. This may reflect genotype-by-diet interactions, as suggested by evidence of its differential expression in response to dietary fatty acid profiles [68].

Due to the limited sample size, we did not investigate pleiotropy among the traits, although they are clearly interrelated and likely share some variants or QTL. Feed intake is integral to FCR and PE, influencing ADG, while PE is theoretically a subset of FCR, with the remaining variance attributed to the conversion of dietary energy into adipose tissue. Consequently, fewer genes may contribute to PE than to FCR, potentially facilitating the detection of PE-associated genes. However, the genetic architectures of PE and FCR likely differ, as suggested by our findings, which identified several plausible regions for PE but none for FCR. We previously reported a genetic correlation of -0.55 ± 0.14 between PE and FCR and -0.53 ± 0.14 between PE and ADFI [10], supporting the assumption that the ability to convert dietary protein into muscle mass (PE) is a component of the ability to convert feed into body mass (see [15] for a more detailed discussion). However, no overlapping regions for these traits were identified in this study. Overlapping regions may exist below detection thresholds, and given the limited number of regions identified, the probability that these regions have an effect on both of these traits will be small. Additionally, none of the highlighted regions for PE on SSC2 and SSC9 matched entries for “feed efficiency” or “feed conversion ratio” in PigQTLdb using the JBrowse tool.

#### Challenges in variant detection for complex and difficult-to-measure traits

The limited ability to detect significant or suggestive associations in this study is likely due to the absence of large-effect SNPs and a relatively small sample size, both key factors for genotype-phenotype associations. Although ADG, FCR, and lean meat content together currently account for only 18 % of the dam line breeding goal (compared to 53 % for the sire line), it is possible that genes with large effects on these traits have been fixed for the favourable alleles during selection history. Even though PE has not yet been included in the breeding goal, this could have been the case due to genetic correlations with the above traits. According to Goddard and Hayes [164], detecting QTL with small effects requires large sample sizes. For this study, with around 1,000 animals and a genomic heritability of 0.42 for PE, only variants with effects >5% could be detected. Most variant effects here ranged from 0 to 1.4%, suggesting a sample size of at least 7,000 would be required to meet the Bonferroni threshold based on all variants.

Achieving the required sample size for PE is challenging due to the complexity of phenotyping this trait and the lack of high-throughput tools for efficient measurement. Models or estimation methods, such as approximating protein content from body weight, are commonly used in nutrition research, but rely on assumptions about an ‘average’ pig [15]. This approach risks masking the individual variation that is critical for estimating breeding values and linking genomic to phenotypic variation. Such inaccuracies and measurement errors can hinder QTL identification [165]. To address this, we used dual X-ray absorptiometry (DXA), which provides highly accurate PE phenotypes based on individual body composition measurements [41]. However, DXA is time consuming, requiring approximately 15 minutes per carcass, which poses a significant challenge for phenotyping large populations.

Our approach to compiling the phenotype data set may have affected the statistical power to detect associated variants. To maximise sample size, we pooled data from different feeding trials with different dietary treatments (e.g. protein content), sex (entire males, females and castrated males) and age at slaughter, resulting in heterogeneity in diet, slaughter weight and sex. The genetic architecture of the traits is likely to vary across these factors. For example, sex differences in PE are well documented, with entire males having higher PE than females and castrated males [41, 43], although it remains unclear whether these differences depend on genotype (genotype-by-sex interactions). Genetic differences in PE across developmental stages are also suggested by genetic correlations below one and partially non-overlapping QTL [25, 26, 43]. In addition, genotype-by-feed interactions may influence results [15]. The polygenic architecture of PE suggests that, while some loci may be specific to certain age classes, sexes, or diets, there are likely also loci associated with PE that are consistent across these conditions, as demonstrated in [25]. Although we accounted for these differences by including covariates in a linear model and using the residuals, this approach may not have fully addressed the issue.

To identify more QTL in GWAS, we used whole-genome sequence data to increase the likelihood of including causal variants, which is expected to improve statistical power. However, as is the case with array data imputed to sequence level, this dramatically increases the number of variants and thus statistical tests, leading to a highly conservative genome-wide significance threshold. Additionally, because variants are not independent due to high LD, the number of tests is effectively inflated. One solution is to base the Bonferroni correction on the number of independent variants rather than the total number. Delpuech et al. [166] used the method by Gao et al. [167] to compute the number of independent SNPs and reduced ∼570K SNPs to 1,690 independent variants, corresponding to a genome-wide threshold of -log10(P) = 4.5. However, with whole-genome data, this approach is computationally very expensive, as it requires computing correlation matrices between all variants per chromosome and then extracting the principal components needed to capture 99.6% of the genotype variability [167]. Instead, we adopted two alternative approaches: permutation testing [56, 57] and LD pruning based on published data on LD decay in commercial pig lines [58]. Permutation testing empirically generates the null distribution of the test statistic, allowing for the determination of an experiment-wide critical threshold. In our case, this resulted in -log_10_(P) values ranging from 7.35 to 7.50. LD pruning, which filters for independent variants (i.e., those below a defined LD threshold) based on typical species- or line-specific LD decay, yielded slightly less conservative thresholds of approximately -log10(P) = 7. However, this method does not rely on the specific data used in the study. For RHM, fewer tests are required as thousands of variants are grouped into windows. We also applied two thresholds, a classical Bonferroni correction (assuming windows are independent, though this may not always hold) and a threshold allowing one false negative per genome scan. These correspond to -log10(p) values of 4.7 (significant) and 3.4 (suggestive).

Another challenge with whole-genome data in RHM is inflated heritability estimates due to high LD of variants within windows. Since LD is unevenly distributed across the genome [58, 168], this can lead to an upward bias in genomic relatedness [169, 170]. We observed this problem in our data, albeit to a minor extent. The genomic heritability estimates for the top 50 RHM windows in our study ranged mostly from 0.8% to 9% (Tables S5–S8), similar to findings in other studies [38, 171, 172]. For PE, heritabilities in the top 50 windows were generally low (0.01 to 0.06), except for one window on SSC9 (37500_42500) with a heritability of 0.12 ± 0.07. Notably, four overlapping windows for ADG on SSC14 had higher heritabilities (suggestive window 637500_642500, h^2^ = 0.15 ± 0.08; 640000_645000, h^2^ = 0.20 ± 0.09; 642500_647500, h^2^ = 0.24 ± 0.1; 645000_650000, h^2^ = 0.18 ± 0.09). Heritabilities for ADFI top 50 windows were also mostly low (0.02 to 0.08) except for two regions on SSC2 (375000_380000 and 380000_385000) with estimates of 0.30 ± 0.16 and 0.27 ± 0.13, respectively. Methods such as reweighting SNPs within a single GRM (LDAK) [173] or stratifying variants into LD and MAF bins across multiple GRMs (GREML-LDMS) [169] can reduce such bias in heritability estimation. A similar issue has been observed in genomic prediction, where higher marker density, such as whole-genome sequence data, only improves accuracy when combined with biological information, as shown in a simulation study [174].

As for genomic prediction, the inclusion of prior biological information may be beneficial for GWAS as well, as it can aid in the detection of associated variants, for instance by using Bayesian hierarchical models [175]. Useful prior information includes previously published QTL, genes and pathways relevant to the trait of interest, supported by detailed functional annotations [176], as well as data from various omics approaches and DNA accessibility studies. Molecular QTL, such as those for the expression of protein-coding genes, non-coding RNAs, enhancers and alternative splicing (e.g. pigGTEx project) [62], are particularly promising. Alternatively, biological information can be assigned to chromosomal regions (e.g. ∼1 Mb) based on their gene content [166, 177]. In general, combining different methods and integrating existing resources can strengthen findings and provide additional evidence for associations that might otherwise fall below the significance threshold. This approach is particularly valuable for traits where phenotyping is challenging and sample sizes are inherently limited, ensuring optimal use of the available data.

## Conclusions

Here, we aimed to associate genomic variants with PE, a trait of particular interest for reducing the environmental impact of pork production through breeding, and other production traits. While GWAS and RHM analyses for PE did not identify significant variants or windows, we identified variants and genomic regions associated with ADFI and ADG, which are genetically correlated with PE. Several genes in these regions are plausible functional candidates for production traits and potentially PE, including those involved in nutrient sensing, the urea cycle, metabolic pathways, in particular IGF1-insulin. Notably, most variants highlighted by GWAS for PE overlapped with top-ranked regions in RHM, providing further evidence for potential associations. While several of the highlighted genes have roles in nitrogen metabolism in cattle, in feed intake and muscle and adipose tissue metabolism in pigs, their specific roles in PE and production traits require further investigation. The role of gene regulation in shaping these traits, in particular the noncoding RNAs and their targets found in this study, should also be addressed in the future. The relatively small sample size, due to the challenges of measuring PE, likely limited the identification of significant variants. Nonetheless, despite the absence of major QTL, the traits showed considerable genomic heritability, suggesting a complex trait architecture with the contribution of numerous small-effect QTL. This genomic information could potentially be leveraged for genomic prediction. However, this requires a large reference population, emphasizing the need for faster phenotyping methods for PE. In summary, despite challenges associated with small sample sizes and difficult-to-measure traits, we identified plausible candidate genes for ADFI and ADG and highlighted potential candidates for PE through overlapping GWAS and RHM results. The development of more sensitive methods and further research into PE remains essential to realise its potential to improve the sustainability of pork production.

## Declarations

### Ethics approval and consent to participate

The experimental procedure was approved by the Office for Food Safety and Veterinary Affairs (2018_30_FR) and all procedures were conducted in accordance with the Ordinance on Animal Protection and the Ordinance on Animal Experimentation.

### Consent for publication

Not applicable.

### Availability of data and material

A preprint of this manuscript has been deposited on bioRxiv (DOI: 10.1101/2023.11.28.568963). Whole genome sequences are available at NCBI Sequence Read Archive (PRJNAxxxxxx; pending). Phenotype data will be made available on Zenodo (DOI:10.5281/zenodo.6985500) after an embargo period of two years.

## Authors’ contributions

EOE curated and analyzed the data, and drafted the manuscript. ALV, AN, and HP participated in the data curation and analysis. CK conceived of the study and participated in its design and coordination. All authors read and approved the final manuscript.

## Acknowledgements

We are grateful to Markus Neuditschko for discussions and comments on the manuscript and to Giuseppe Bee for discussions. We thank Dorothea Lindtke for help with bioinformatics and discussion about the significance thresholds, Valentina Riggio for advice concerning the RHM analysis, and Paolo Silacci and his team at Agroscope’s Animal Biology Group for DNA extraction from blood.

## Footnotes

1 Fütterungsempfehlungen und Nährwerttabellen für Schweine (Feeding recommendations and nutrient tables for pigs). Agroscope, Posieux, Switzerland. Retrieved 31 January f from https://www.agroscope.admin.ch/agroscope/fr/home/services/soutien/aliments-pour-animaux/apports-alimentaires-recommandes-pour-les-porcs.html

2 https://www.ncbi.nlm.nih.gov/gdv?org=sus-scrofa&group=artiodactyl^a^

3 https://piggtex.ipiginc.com/

